# *In Silico* Optimisation of Regenerative Cell Therapy in the Infarcted Human Ventricles to Mitigate Arrhythmic Burden

**DOI:** 10.1101/2025.10.03.680241

**Authors:** Leto Luana Riebel, Zhinuo Jenny Wang, Xin Zhou, Lucas Arantes Berg, Cristian Trovato, Blanca Rodriguez

**Affiliations:** Department of Computer Science, University of Oxford, Oxford, UK; Systems Medicine, Clinical Pharmacology & Safety Science, R&D, AstraZeneca, Cambridge, UK

## Abstract

Myocardial infarction remains a frequent cause of heart failure and mortality. Cell therapy has been shown promising in pre-clinical trials to regenerate the damaged tissue, but delivered cells may beat spontaneously and produce arrhythmias in the ventricles, particularly in the first weeks after delivery, which hinders clinical application. Previous studies have proposed ionic targets to supress the cells’ automaticity but, so far, the effects of such treatments on the cells’ calcium dynamics, as a key driver of contractile function, have been insufficiently evaluated. Furthermore, effective strategies are needed that can alleviate the injected cells’ pro-arrhythmic action potential phenotypes.

The goal of our study was to identify mechanisms to mitigate arrhythmic pathways following cell delivery in the chronically infarcted human ventricles using multiscale modelling and simulation. First, we demonstrate credibility by simulating experimentally observed transient automaticity-induced ventricular tachycardia arrhythmias with a frequency of up to 140 beats per minute at two weeks post cell injection in three different infarct geometries. Next, our simulations show how the timeframe during which re-entry was inducible increases 1) from before to after cell delivery at day 0 from 0 to 640 ms, 60 to 100 ms, and 60 to 760 ms in the small, medium, and large scar, respectively, and 2) from day 0 to day 14 after virtual cell injection in the large scar by 175%. Finally, we show that a combination of blocking the funny current and upregulating the inward rectifier potassium current, the sodium potassium pump, and the rapid delayed outward rectifier potassium current can reduce both automaticity-induced and re-entrant arrhythmias while maximising calcium amplitudes.

In conclusion, our simulations show that not only automaticity-induced but also re-entrant arrhythmias increase as injected cells mature in the ventricles and depend on the scar size. Furthermore, through modelling and simulation, we identify anti-arrhythmic strategies to improve therapy safety while maximising efficacy.

## Introduction

Myocardial infarction (MI) and its long-term consequences, particularly heart failure, remain amongst the leading causes of death worldwide. Since adult human hearts do not sufficiently regenerate on their own, myocardial damage is largely irreversible. To ease the global burden of heart failure and improve cardiac function post-MI, delivery of stem cells has been explored (Jebran et al., 2025; Liu et al., 2018; Shiba et al., 2016). Whilst pre-clinical studies have shown promising effects, ventricular arrhythmias, particularly during the first weeks after cell injection, have been observed and are suspected to arise from the immature cells’ spontaneous beating (Chong et al., 2014; Liu et al., 2018; Marchiano et al., 2023; Romagnuolo et al., 2019).

Previous studies have identified upregulation of the inward rectifier K^+^ current (I_K1_) and suppression of the funny current (I_f_) as crucial anti-automaticity targets (Goversen et al., 2018; Kim et al., 2015; Verkerk & Wilders, 2023). One recent study found a combination of these together with a block of the T-type Ca^2+^ current (I_CaT_) and the sodium calcium exchanger current (I_NaCa_) necessary to suppress spontaneous beating (Marchiano et al., 2023), highlighting a complex mechanism behind the cells’ automaticity. Whilst these strategies offer a vital way to reduce arrhythmias, it is unclear how they might impact the cells’ calcium dynamics and, consequently, their efficacy. In addition to automaticity-induced arrhythmias, previous studies have highlighted a variety of re-entrant pathways caused by the cells’ immature tissue conductivity and action potential (AP) phenotype (Fassina et al., 2023; Riebel et al., 2024; Yu et al., 2022). To enable safe clinical application of cell therapy post-MI, effective anti-automaticity as well as anti-re-entry targets need to be identified.

The development and testing of new treatments, whether they reach the clinic or not, is expensive (Sertkaya et al., 2024). Multiscale human modelling and simulation offers a power technology to augment the drug discovery and development process by identifying safety risks early on (Passini et al., 2021; Viceconti, Emili, et al., 2021). Furthermore, simulation studies present an efficient tool to design anti-arrhythmic strategies that improve therapy success (Dasí et al., 2024; Rosales et al., 2024).

In this study, we harness the powers of human-based modelling and simulation to identify and optimise ionic targets that reduce both automaticity-induced and re-entrant arrhythmias after cell delivery, as a post-MI therapy. For this, we conduct computer simulations from ionic current dynamics to the electrocardiogram (ECG) to evaluate pro-arrhythmic mechanisms following cell delivery in in the infarcted human ventricles including the Purkinje system (Riebel et al., 2024). We based our modelling and simulation of maturing injected cells on a substantial experimental dataset including experimentally-reported gene expression data from Marchiano et al. (2023) of how injected cells mature *in vivo*. We then set out to determine 1) the risk and mechanisms of automaticity-induced arrhythmias at day 0 (D0) and day 14 (D14) after cell injection, and how it is modulated by scar size, 2) the re-entry risk before and at D0 and D14 after delivery, 3) novel ionic targets that supress spontaneous beats whilst maintaining physiological calcium transients (CaT), and 4) a combination of ionic targets to optimise therapy safety by reducing automaticity-induced as well as re-entrant arrhythmias.

## Results

### Establishing credibility: Simulating automaticity-induced arrhythmias two weeks after cell delivery

As expected from experimental observations (Chong et al., 2014; Marchiano et al., 2023; Romagnuolo et al., 2019) and in line with our previous study (Riebel et al., 2024), human stem cell-derived cardiomyocytes (hPSC-CMs) immediately after delivery (D0) did not produce spontaneous beats in the ventricles (see Figure 1A). Two weeks following cell delivery (D14), hPSC-CMs caused ectopy in the ventricles with 42 beats per minute (bpm). Similarly, a rapid hPSC-CM phenotype (see Methods for detail), beat 46 times per minute at D0. However, both were overridden by 1 Hz sinus pacing (see the lack of automaticity in Figure 1B & C).

**Figure 1.**
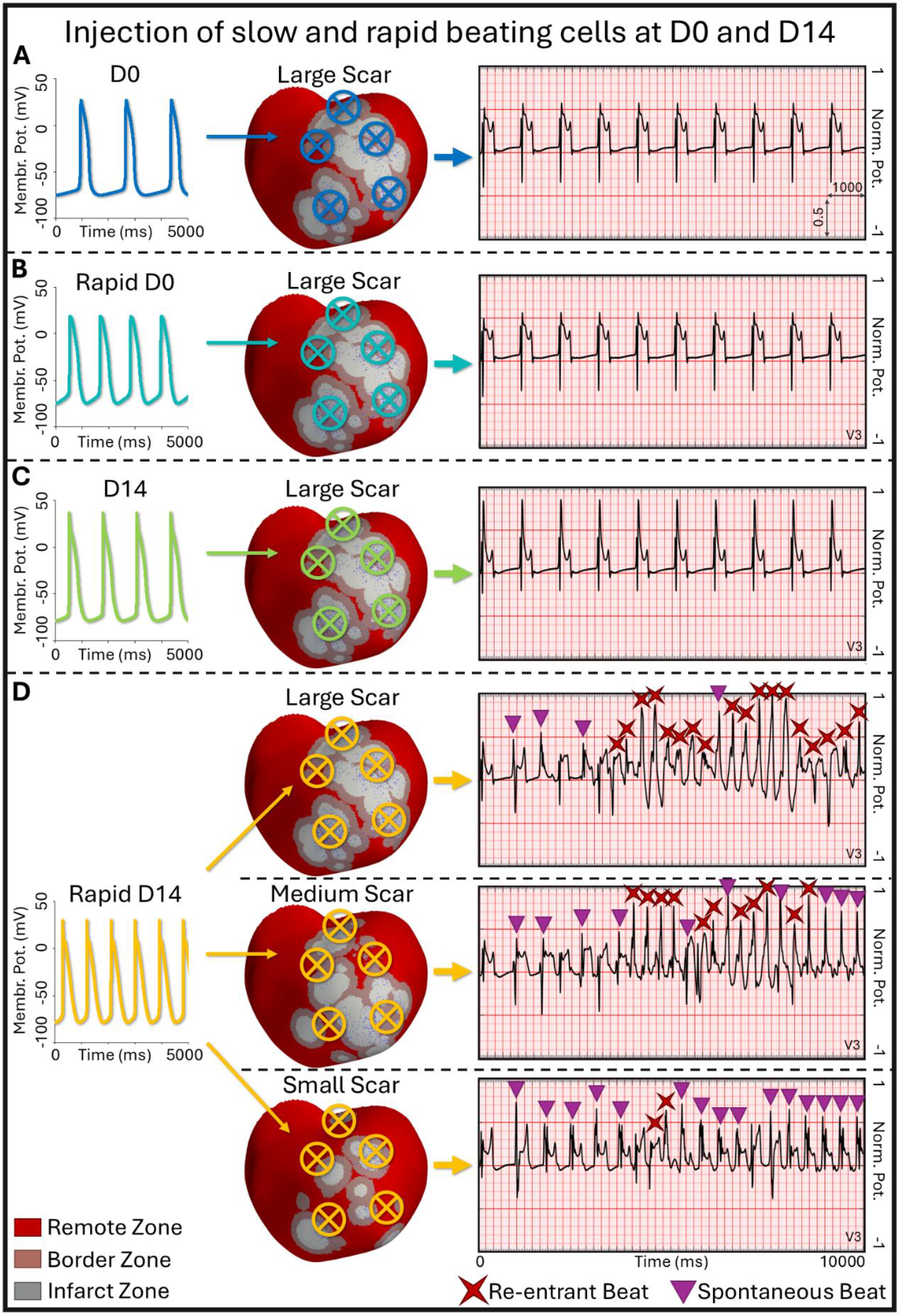
Automaticity-induced arrhythmia burden for virtual injection of baseline and rapid hPSC-CMs at D0 and D14 after delivery in three different human biventricular models of chronic infarction with Purkinje. Highlighted with crosses in the biventricular mesh are five out of six virtual hPSC-CM injection locations (the remaining one is located in the septum). Also see Video S1.

At D14 post-delivery and as shown in Figure 1D, the rapid phenotype, which beats 66 times per minute in single cell, produced spontaneous depolarisations in all three investigated scars. Spontaneous beating increased over time and facilitated VT-like arrhythmias with intermittent re-entrant wavefronts in all three scars. As highlighted in the annotated ECGs of Figure 1D, the proportion of re-entrant wavefronts compared to spontaneous depolarisations decreased with decreasing scar size (from 82% in the large scar over 52% in the medium scar to 12% in the small scar). While re-entry dominated over spontaneous activity in the large scar, automaticity appeared to reach a state-state towards the end of the 10 s simulations in the medium and small scar (see Video S1). Spontaneous depolarisations occurred up to every 420 ms in the ventricles corresponding to a VT frequency of approximately 140 bpm, similar to pre-clinical observations (Marchiano et al., 2023). Spontaneous hPSC-CM beating was faster in the ventricles than in single cell (140 versus 66 bpm); as discussed in our previous work (Riebel et al., 2024), hPSC-CM automaticity in the ventricles is favoured by depolarised chronic infarct scars and may further increase under fast pacing, which here was facilitated through re-entrant rotors (see ECGs in Figure 1D).

### Quantifying re-entry risk: Arrhythmic burden increases from before to after cell delivery and from day 0 to day 14

We next set out to quantify the risk of re-entry after cell delivery. As shown in Figure 2, using an S1-S2 stimulus protocol, we observed few re-entries before virtual cell injection. In the small scar, which contained spread out scar islands, no re-entries were observed. In the medium and large scar each, two non-sustained and one sustained re-entry occurred. All re-entries in the large scar and two out of three in the medium scar were induced by ectopic stimulus locations close to the apex and followed a regular pattern. As the scar cores were unexcitable and only slowly conducting, the wavefront travelled through isthmuses between the dense scar cores and re-entered at the site of the ectopic stimulus. The third re-entry in the medium scar was facilitated through a Purkinje-myocyte junction (PMJ) on the anterior left ventricular (LV) wall on the edge between infarct (IZ) and border zone (BZ).

**Figure 2.**
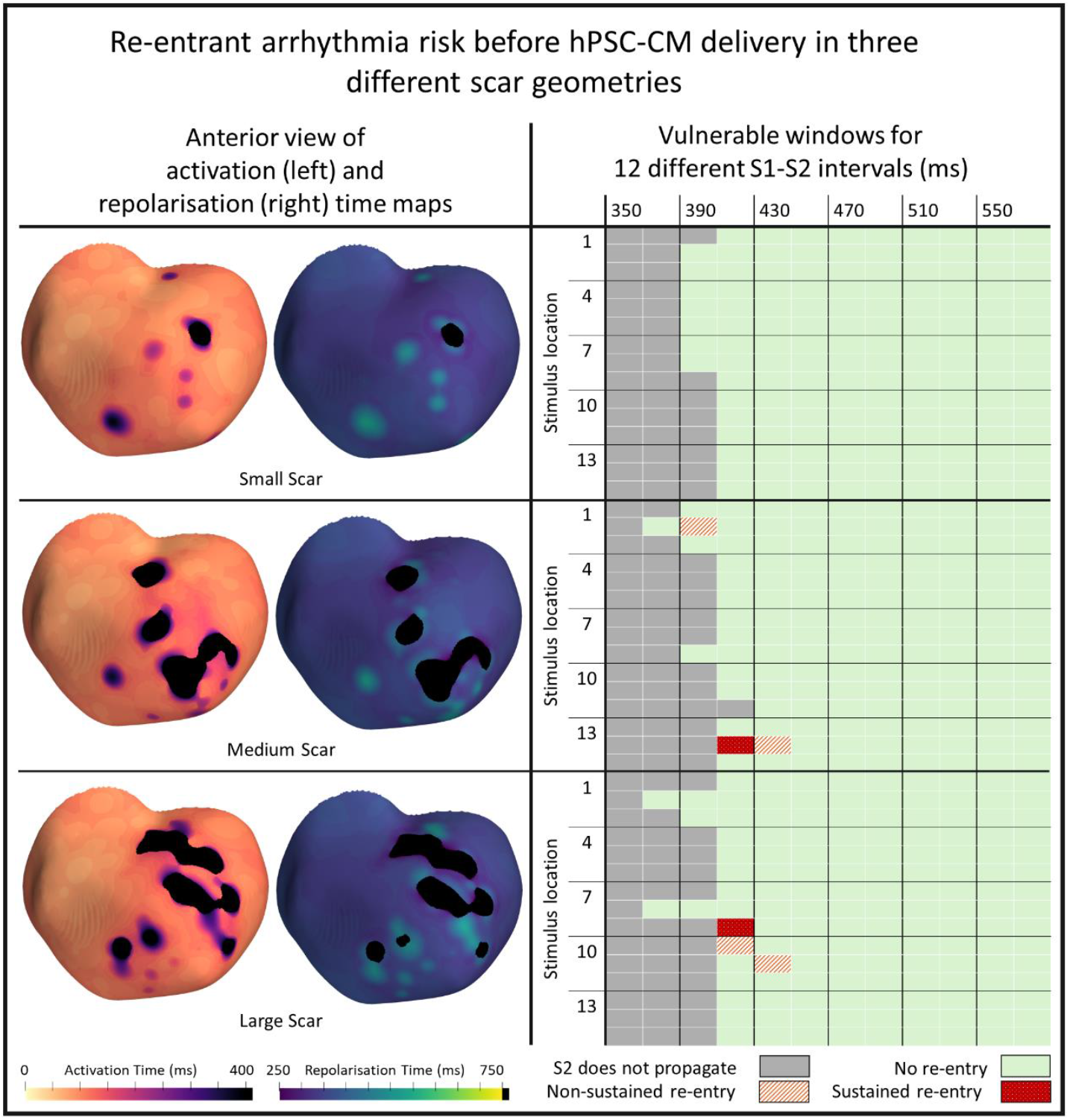
Re-entry risk in a human biventricular model with three different scar geometries before virtual hPSC-CM delivery. Left: Activation and repolarisation time maps. Right: VWs for different S1-S2 intervals and S2 stimulus locations. Note, ectopic stimulus locations are different in each scar and were ordered from most anterior-basal to most inferior-apical.

At D0 after delivery and as shown in Figure 3, the vulnerable windows (VWs) increased in all three investigated scars: from 0 to 640 ms in the small scar, from 60 to 100 ms in the medium scar, and from 60 to 760 ms in the large scar (see Figure 2 and Figure 3). While re-entry risk increased in all three scar sizes, it did not increase with increasing scar size (i.e, re-entry risk was highest in the large scar but lowest in the medium scar). However, re-entry sustainability was lowest in the small scar (3%, i.e., 1 out of 32 re-entries after cell delivery) and substantially higher in the medium (33% before and 80% after delivery) and large scar (33% before and 76% after delivery). Interestingly, all re-entries in the small scar occurred through one of two PMJs in the centre of two circular scar islands (see Video S2). These allowed for figure-of-eight re-entries and demonstrate the crucial role of scar morphology in arrhythmogenesis. In the medium and large scar, re-entries occurred through tissue as well as PMJs. While in the medium scar 80% of re-entries involved the same large apical scar core (visible as not fully repolarising in the repolarisation map of Figure 3), no area within the large scar appeared particularly pro-arrhythmic. As highlighted when comparing the activation time maps in Figure 2 and Figure 3, cell delivery improved conductivity and excitability of the IZ and allowed for wavefronts to travel through infarct cores, which before cell delivery were inaccessible (see Video S3).

**Figure 3.**
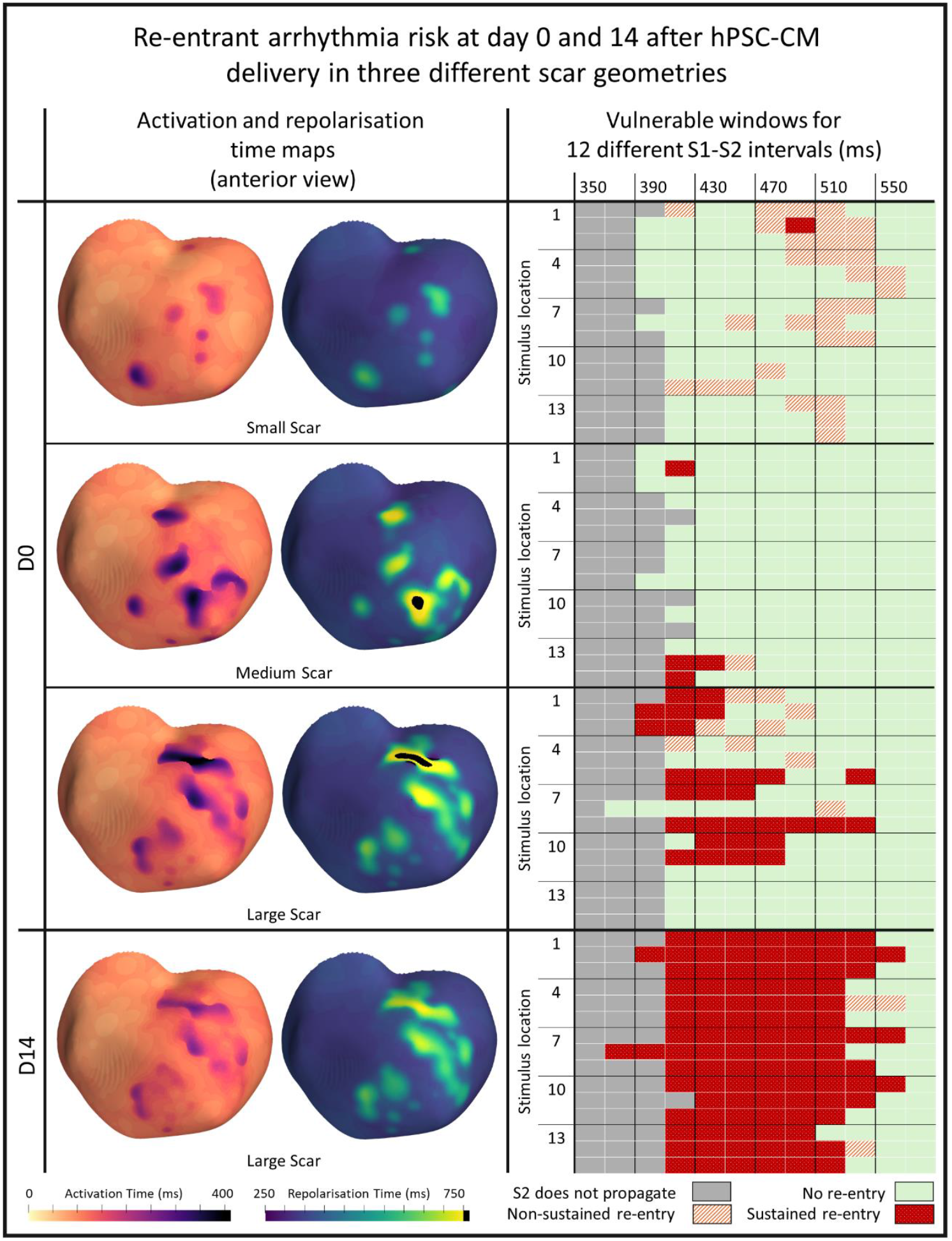
Re-entry risk in a human biventricular model with three different scar geometries at D0 and D14 after virtual hPSC-CM delivery. Left: Activation and repolarisation time maps. Right: VWs for different S1-S2 intervals and S2 stimulus locations. Note, ectopic stimulus locations are different in each scar and were ordered from most anterior-basal to most inferior-apical. Also see Videos S2 and S3.

Next, we investigated the re-entry risk at D14 post-delivery in the large scar (as this was the most pro-arrhythmic scenario at D0); the VW increased from 760 to 2,080 ms, as shown in the bottom of Figure 3. 97% (101 out of 104) of those D14 re-entries sustained for at least two re-entry cycles. As highlighted by the repolarisation time maps in Figure 3, D14 cells further improved conductivity within the IZ. This was due to an increase of conduction velocity (CV) from 10 cm/s at D0 to 14.5 cm/s at D14 – as conductivities were maintained, this was caused by changes in ionic current conductances (see Methods section). The improved conductance within the IZ enabled new paths for the re-entrant wavefront; slow transmural conduction pathways emerged as particularly pro-arrhythmic at two weeks post injection.

To summarise, compared to the adult ventricular cardiomyocytes, the delivered cells’ prolonged AP duration at 90% repolarisation (APD_90_) promoted ventricular repolarisation gradients, refractoriness, and pro-arrhythmic unidirectional conduction block. Whilst increased CV within the chronic scar after cell delivery reduced activation time gradients in the ventricles, it also enabled new re-entry pathways transmurally and through the scar core, particularly at D14 after delivery.

### Target discovery: Regulating I_NaK_ to supress automaticity-induced arrhythmias

To identify ionic targets that suppress hPSC-CM automaticity, we carried out a cellular sensitivity analysis. Our results, shown in Figure S1, identified the spontaneous beating frequency to be most sensitive to I_NaK_, with both a 45% increase as well as a 30% decrease in its maximum value (p_NaK_) abolishing automaticity at D0. As shown in Figure 4A, reducing p_NaK_ caused intracellular Na^+^ accumulation leading to an increase in the reverse mode of I_NaCa_ and, ultimately, intracellular Ca^2+^ accumulation and APD shortening, as discussed in other studies (Bueno-Orovio et al., 2014; Faber & Rudy, 2000). Counterintuitively and as depicted in Figure S2A, intracellular Na^+^ accumulation over time resulted in an ultimately increased I_NaK_, even though p_NaK_ was reduced. Further investigation in our model also showed an increase in the SR Ca^2+^ concentration. Under unpaced conditions, a corresponding impaired recovery of the RyR-sensitive release current’s (I_rel_) inactivation gate (model parameter RyRc) leading to a reduced Ca^2+^-induced Ca^2+^ release through I_rel_ resulted in a lack of spontaneous activity. While 1 Hz pacing caused physiological APs at D0 (see Figure S2A), SR Ca^2+^ overload occurred and only diminished CaTs and APs with a short duration could be produced at D14 (see Figure S3A). As such AP heterogeneities may be pro-arrhythmic, we discarded p_NaK_ downregulation and considered its upregulation further.

**Figure 4.**
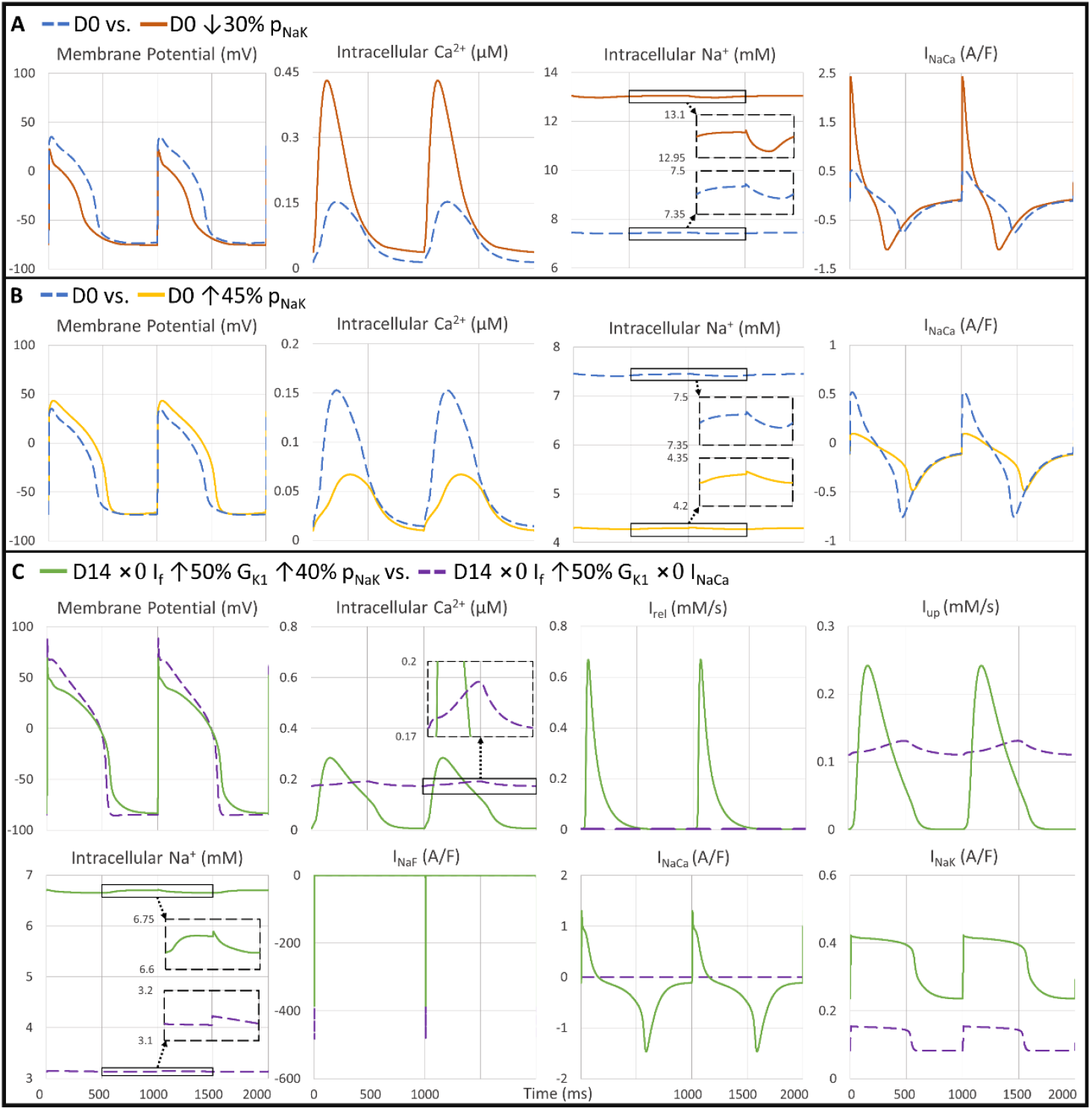
APs and ionic current traces of different hPSC-CM phenotypes in single cell after 1,000 paced 1 Hz beats. A) hPSC-CMs at D0 with and without downregulating pNaK. B) hPSC-CMs at D0 with and without upregulating p_NaK_. C) hPSC-CMs at D14 with full I_f_ block, upregulation of G_K1_ and G_Kr_, and either full I_NaCa_ block (Marchiano et al., 2023) or p_NaK_ upregulation.

Upregulation of p_NaK_ suppressed spontaneous beats through the opposite effect, i.e., intracellular Na^+^ and the reverse mode of I_NaCa_ were reduced therefore leading to lower intracellular and SR Ca^2+^ concentrations impairing the Ca^2+^ clock (see Figure 4B and S2B). At D14, p_NaK_ upregulation on its own was not sufficient to abolish automaticity and hence we combined it with a full block of I_f_ and upregulation of I_K1_’s conductance (G_K1_), as suggested in the literature (Goversen et al., 2018; Marchiano et al., 2023). For a 50% increase in G_K1_, a 40% and 65% increase in p_NaK_ was enough to stop spontaneous beating in the baseline and rapid phenotypes, respectively. Pacing produced physiological APs and CaTs at D0 and D14, as shown in Figures S4 and S3B, respectively. We further compared our suggested p_NaK_ upregulation with the state-of-the-art approach of blocking I_NaCa_ to inhibit stem cell automaticity (Marchiano et al., 2023). As depicted in Figure 4C, this comparison showed that together with increased G_K1_ and blocked I_f,_ I_NaCa_ block caused intracellular Ca^2+^ accumulation and small CaT amplitudes (CaT_Amp_). We identified the main mechanism behind this as a substantially reduced I_rel_, similar to under p_NaK_ downregulation. A block in I_NaCa_, which is the main mechanism for Ca^2+^ extrusion, caused accumulation of intracellular but also SR Ca^2+^. This led to insufficient relaxation of I_rel_’s activation and inactivation gates (RyRo and RyRc model parameters). In comparison, even though the CaT_Amp_ was reduced compared to the baseline models with no ionic current modifications, p_NaK_ upregulation produced more physiological CaTs at both D0 and D14 (see Figure S4 and Figure 4C, respectively).

### Target optimisation: Combining I_f_ block with G_K1_, p_NaK_, and G_Kr_ upregulation to reduce automaticity-induced as well as re-entrant arrhythmias

Next, we investigated how to combine the identified anti-automaticity targets, i.e., blocking I_f_ and upregulating G_K1_ and p_NaK_, with anti-re-entry targets. Our sensitivity analysis, shown in Figure S1, as well as previous studies (Fassina et al., 2023; Riebel et al., 2021), highlighted the rapid delayed outward rectifier K^+^ current (I_Kr_) as a promising target to modulate the increased stem cell APD towards adult values. Hence, we created an automated algorithm aiming to determine scaling factors for upregulating G_K1_, p_NaK_, and I_Kr_’s conductance (G_Kr_) in addition to full I_f_ block. As described in the Methods, this algorithm aimed to optimise both CaT_Amp_ and APD_90_ by maximising an optimisation index that considers both values. Using a target APD_90_ of 330 ms (approximately ^3/4^ of the paced D0 APD_90_), scaling G_K1_ by 1.5, p_NaK_ by 1.75, and G_Kr_ by 3 returned the highest optimisation index (see Figure S5). At D0, this optimisation decreased hPSC-CM APD_90_ to 284 ms and increased CV from 10 cm/s to 12.5 cm/s. However, in biventricular simulations, these ionic scalings reduced the VW only marginally from 760 to 680 ms at D0, as shown in Figure 5A. Increased CV improved activation of the scar (see repolarisation time maps in Figure 3 and Figure 5A) and, similarly to the unoptimized phenotype at D14, enabled new re-entrant pathways.

**Figure 5.**
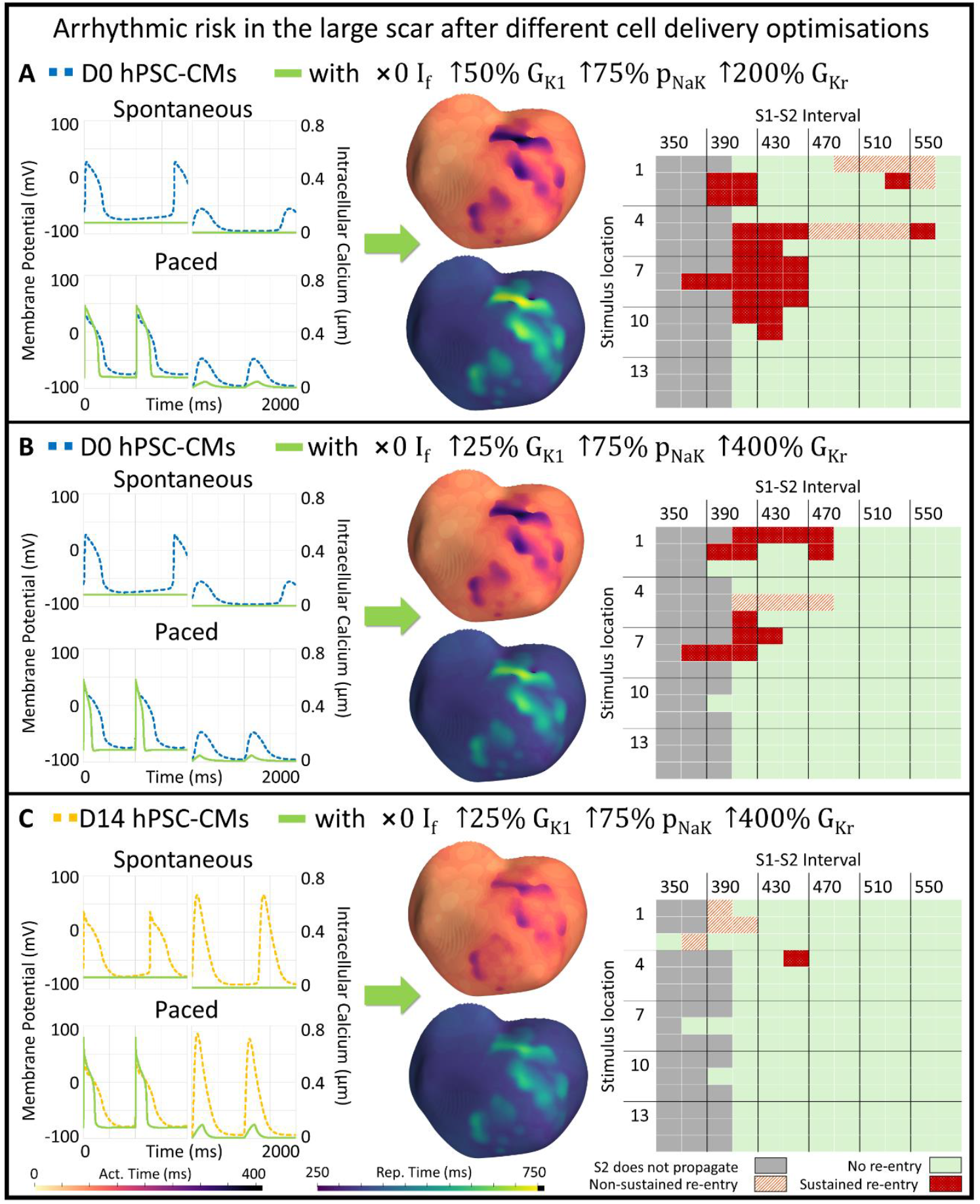
Re-entry risk at D0 and D14 after virtual delivery of optimised hPSC-CMs into a human biventricular chronic post-MI model. Left: APs and CaTs. Middle: Activation and repolarisation time maps. Right: VWs for different S1-S2 intervals and S2 stimulus locations. In panel C, the ectopic stimulus for location 3 and S1-S2 interval of 330 ms (not shown in the table) did not propagate. Note, ectopic stimulus locations were ordered from most anterior-basal to most inferior-apical. Also see Video S4.

As repolarisation gradients were not sufficiently reduced, we proceeded to further shorten the target APD_90_, by about half, to 170 ms (see results in Figure S6). As shown in Figure 5B & C, after upregulating G_K1_ by 1.25, p_NaK_ by 1.75, and G_Kr_ by 5 (in addition to full I_f_ block), VWs decreased from 760 to 340 ms at D0 and from 2,080 to 100 ms at D14. Decreased arrhythmogenicity was the result of an increased CV and reduced APD within the scar, which homogenised both activation and repolarisation gradients, respectively (see Figure 5B & C and Video S4). Optimisation increased hPSC-CM CV by 25% and 24% at D0 and D14, respectively. Since conductivities were kept constant, this was caused by blocking I_f_ as well as upregulating G_K1_, p_NaK_, and G_Kr_, which have all been shown to improve AP upstroke velocity under paced conditions (Riebel et al., 2021). As shown in the CaTs in Figure 5B & C, while our algorithm aimed to maximise the CaT_Amp_, it was still substantially lower than before optimisation. This was largely caused by a block in I_f_, which on its own reduced the CaT_Amp_ by approximately 40%.

## Discussion

In this study we present a novel *in silico* multiscale human electrophysiological framework to optimise new treatments such as regenerative cell therapy. First, in a human biventricular model with three different post-MI scars, we showed that while cells remained quiescent immediately after delivery, rapid cells produced automaticity two weeks later, causing monomorphic ectopy-driven VT in the small and medium scar, and polymorphic re-entry driven VT in the large scar. Next, in the absence of spontaneous beats, we quantified the re-entry risk, which increased from before to after cell delivery and from D0 to D14. Finally, we demonstrated how our framework enables identifying and optimising anti-arrhythmic ionic target combinations – with a block of I_f_ and upscaling of maximum I_NaK_ as well as the conductances of I_K1_ and I_Kr_ emerging as a novel strategy to suppress spontaneous beating and reduce re-entrant arrhythmia susceptibility while maximising CaT_Amp_.

Our *in silico* framework of MI and including the Purkinje network was constructed, calibrated, and evaluated using human-based experimental and clinical evidence and incorporates extensive biophysical knowledge gained throughout the development of modelling and simulation technologies (Camps et al., 2024; Mincholé et al., 2019; Tomek et al., 2019; Zhou et al., 2024). As described in the Methods section, credibility of our human-based simulation framework is provided by addressing verification, validation, and uncertainty quantification principles (Musuamba et al., 2021; Viceconti, Pappalardo, et al., 2021) in the present and previous studies and specifically in Riebel et al. (2024).

Our simulations showed that rapid phenotypes may beat spontaneously in the ventricles two weeks after cell delivery and facilitate VT-like arrhythmias with a frequency of up to 140 bpm, as observed experimentally (Marchiano et al., 2023). Maturation of hPSC-CMs has been investigated also in other *in silico* studies; Paci et al. (2012) used experimental evidence to construct early and late maturation cells. Similar to our study, their late-stage model had increased I_K1_, I_to_, I_NaF_, I_NaCa_, and I_CaL_, but, contrary to our approach, decreased I_f_ and I_Kr_ (Paci et al., 2012). According to Marchiano et al. (2023), while the genes encoding I_f_ and I_Kr_ increase initially, possibly contributing to the surge of arrhythmias two weeks after injection, they ultimately decrease by day 84. It should be noted, that Marchiano et al. (2023) investigated gene expression of hPSC-CMs that had matured for 18-20 days *in vitro* before delivery. Furthermore, as we derived *in silico* D0 and D14 phenotypes from experimental gene expression data (Marchiano et al., 2023), which may not directly infer ionic current conductance, future work may create D0 and D14 hPSC-CM populations to assess this uncertainty.

Next, we utilised the mechanistic nature of modelling and simulation to suggest anti-arrhythmic strategies, with the Na^+^-K^+^ pump emerging as a novel target. Interestingly, experimental studies have suggested that I_NaK_ may increase as hPSC-CMs mature (Otsu et al., 2005), which could contribute to the reduction of automaticity after three weeks post injection and support our findings. The effects of I_NaK_ regulation on spontaneous beating, although to a lesser extent, have also been highlighted in a sensitivity analysis by Paci et al. (2012). Here, in addition to p_NaK_ upregulation, we suggest a full block of I_f_ and upregulation of G_K1_ to suppress spontaneous beating. Whilst our results indicate that for some hPSC-CM phenotypes targeting p_NaK_ may be sufficient, modulating also I_f_ and I_K1_ may be required in others. I_f_ and I_K1_ have long been known to impact the immaturity and automaticity of hPSC-CMs (Goversen et al., 2018; Marchiano et al., 2023; Verkerk & Wilders, 2023). Whilst in our D0 (but not D14) model, I_f_ block and sufficient G_K1_ upregulation stopped automaticity, it has been suggested in other work that this may be insufficient due to the Ca^2+^ clock acting as a second depolarising mechanism (Kim et al., 2015; Marchiano et al., 2023).

Our next goal was to systematically investigate ionic current combinations that minimise arrhythmic risk whilst allowing for physiological CaTs. Current state-of-the-art agrees to block I_f_ and upregulate I_K1_ but instead of targeting I_NaK_ further suggests knock-out of I_CaT_ and I_NaCa_ (Marchiano et al., 2023). As we have shown in our comparison of targeting I_NaK_ versus I_NaCa_, a full block of I_NaCa_ may result in a reduced CaT_Amp_ and Ca^2+^ accumulation, which, although to a lesser extent, has been suggested also in the supplemental material of Marchiano et al. (2023). However, as it is a main source of Ca^2+^ extrusion (Ottolia et al., 2013), reduced I_NaCa_ is generally associated with an increase in CaT_Amp_ despite decreased I_CaL_ (Ismaili et al., 2022; Ozdemir et al., 2008). Nevertheless, full I_NaCa_ knock-out can facilitate intracellular and SR Ca^2+^ accumulation and may ultimately hinder effective contractility (Henderson et al., 2004; Koivumäki et al., 2018, Figure 3; Koushik et al., 2001). Modulating I_NaK_, as we have done here, will ultimately impact also I_NaCa_ as the two currents rely on each other’s regulation of Na^+^ concentrations. Interestingly, amiodarone has been suggested as an anti-arrhythmic strategy in pre-clinical trials of cell delivery (Nakamura et al., 2021) and is known to impact both, I_NaCa_ and I_NaK_ (Gray et al., 1998; Watanabe & Kimura, 2000).

Finally, to reduce not only automaticity but also re-entries, which we showed to increase from D0 to D14, we further suggest an increase of G_Kr_ (in addition to blocking I_f_ and upregulating

G_K1_ and p_NaK_). Targeting I_Kr_ to make hPSC-CM APs more adult-like has been put forward also in other *in silico* studies, which further identified I_NaF_ as a potential target (Fassina et al., 2022, 2023). Besides electrophysiological targets, *in silico* investigations have also highlighted the crucial role of hPSC-CM and scar conductivity on arrhythmic risk (Fassina et al., 2023; Gibbs et al., 2023; Rosales et al., 2024; Yu et al., 2022). Here, we used an hPSC-CM CV of 10 cm/s at D0, which increased to 14.5 cm/s at D14. While the experimental study we consulted to calibrate D14 CV shows a lower D0 CV, it investigated cell fibres without transplanting them (Hansen et al., 2018).

While our suggested optimised ionic target combination includes a 25% increase in G_K1_, our results show that with concurrent I_f_ block and sufficient p_NaK_ and G_Kr_ upregulation, targeting G_K1_ may not be necessary to stop spontaneous beats. Ultimately, optimal scaling factors will vary between cell types, maturation protocols, and the patient, e.g., patients with a longer APD may require the hPSC-CMs’ APD to be shortened less. The computational pipeline we have presented here allows for fast and efficient optimisation and testing of new treatments for a range of different cell types, delivery modalities, and patient substrates.

## Methods

### Multiscale electrophysiological modelling and simulation of the chronically infarcted human ventricles with Purkinje

Electrophysiological activity of the infarcted human ventricles was simulated from subcellular dynamics to the ECG as in (Riebel et al., 2024) (Figure 6). Briefly, a healthy human biventricular geometry was reconstructed from MRI by Mincholé et al. (2019). Next, fibre orientations were introduced as described in Doste et al. (2019) and CV in fibre orientation calibrated to 65 cm/s, as observed clinically (Taggart et al., 2000). Sheet and sheet-normal CV was inferred (38 and 47 cm/s, respectively) and a Purkinje tree created to reproduce the patient’s QRS complex (Camps et al., 2024), see Figure 6A.

**Figure 6.**
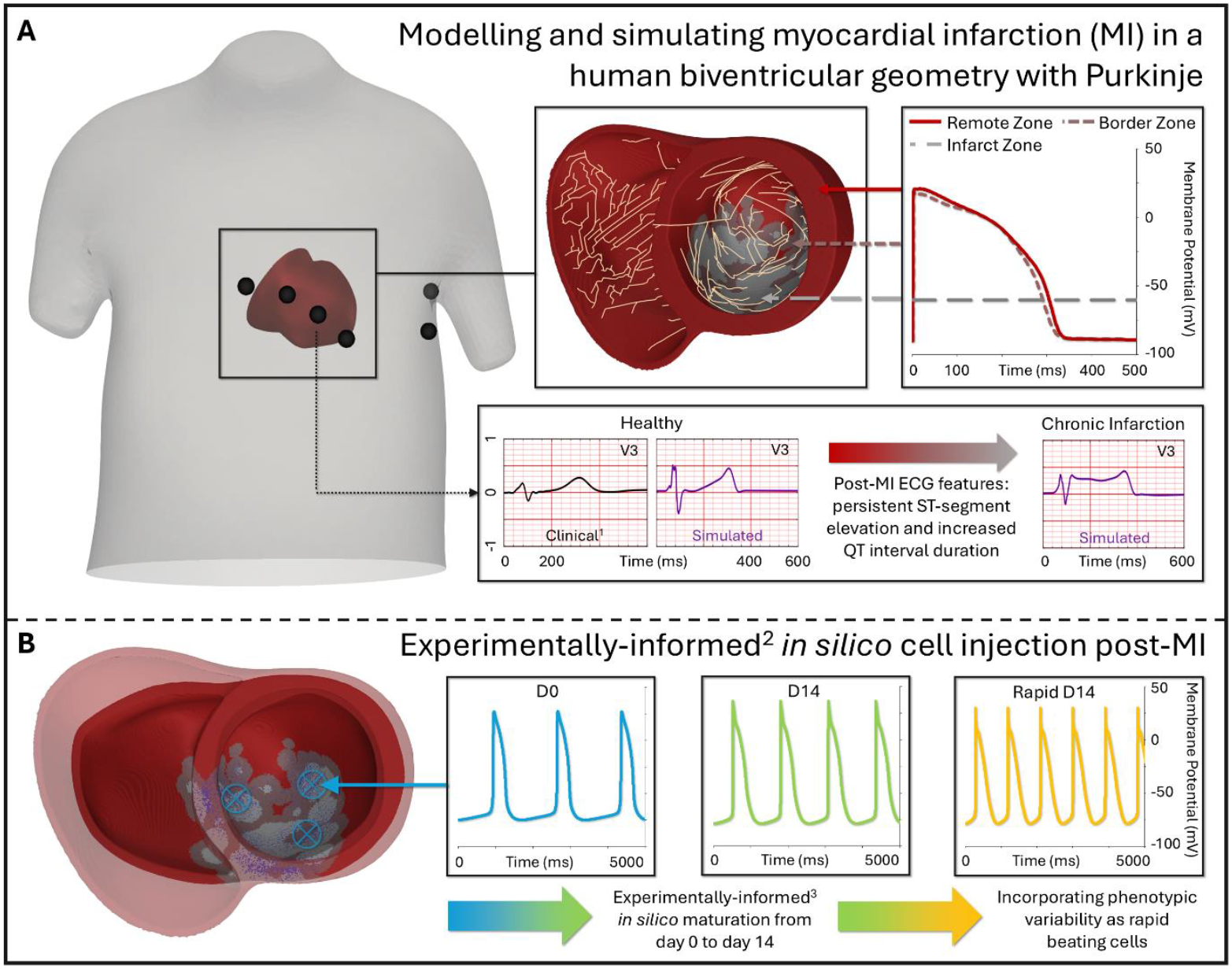
Multiscale modelling and simulation of human ventricular electrophysiology in A) chronic MI and B) cell therapy post-MI using a D0, D14, and rapid beating hPSC-CM phenotype. ^1^Biventricular geometry and ECG recordings from Mincholé et al. (2019). ^2^Spatial distribution of hPSC-CMs was informed by Poch et al. (2022). ^3^Maturation from D0 to D14 was simulated based on in vivo gene expression data from Marchiano et al. (2023).

Membrane kinetics were simulated with the state-of-the-art human ventricular cardiomyocyte and Purkinje cell ToR-ORd and Trovato models, respectively (Tomek et al., 2019; Trovato et al., 2020). Based on clinical observations (Boukens et al., 2015; Franz et al., 1987), biventricular electrophysiological gradients of 35 ms transmurally and 10 ms from apex to base were introduced by scaling key ionic current conductances (summarised in Table S1).

To model MI, three physiological scar geometries were introduced into the healthy geometry using an established algorithm (Cardone-Noott et al., 2014; Hill et al., 2016). Briefly, six landmarks were selected to model the path of the left anterior descending artery along the anterior and septal walls and the apex. In line with clinically reported scar sizes (Reindl et al., 2020; Spath et al., 2021), resulting IZ and BZ composed 14%, 23%, and 27% of the ventricles (referred to as the small, medium, and large scar).

For remote zone (RZ) and BZ electrophysiology, key ion channel conductances and time constants were scaled based on experimental observations as described by Zhou et al. (2024) and summarised in Table S2; resulting APs are shown in Figure 6A. The scar was modelled as electrically inactive but conducting (Rog-Zielinska et al., 2016). Hence, the membrane potential was initialised to −60 mV and allowed to change only through terms of diffusivity (Ringenberg et al., 2014). Electrophysiological changes caused RZ CV to decrease to 49, 27, and 34 cm/s in fibre, sheet, and sheet-normal direction, respectively. Compared to the RZ, conductivities were calibrated to achieve an approximately 60% and 80% reduced CV in the BZ and IZ, respectively (see Table S3), based on clinical observations (Aronis et al., 2020; Jamil-Copley et al., 2015). The resulting simulated ECG traces were compared to clinical data of post-MI subjects in our previous study (Riebel et al., 2024) and reproduced persistent ST-segment elevation, QT-interval prolongation, and normalised T-waves (see Figure S7A).

3D monodomain simulations were performed using the finite volume GPU-enabled open-source solver MonoAlg3D (Berg et al., 2025; Sachetto Oliveira et al., 2018) with parameters listed in Table S4.

### Simulating injection of day 0, day 14, and rapid stem cell-derived cardiomyocytes

As shown in Figure 6B, to investigate transient ectopy-induced arrhythmias, we constructed a D14 hPSC-CM phenotype by modifying ionic conductances of the baseline Paci2020 model (Paci et al., 2020), which we consider the D0 phenotype. For the D14 model, experimentally reported *in vivo* gene expression data (Marchiano et al., 2023) was consulted to derive ion channel scaling factors. As data to exactly relate ion channel conductance from gene expression is lacking, to qualitatively capture the cell’s maturation, we manually estimated the relative change in gene expression (Marchiano et al., 2023) and multiplied it by 0.25 to give a scaling factor for the corresponding ion channel conductance, as summarised in Table S5. This scaling factor of 0.25 accounts for the differences between the transcription and the translation level, considering factors such as translation efficiency, protein stability, and regulatory feedback (Stevens & Brown, 2013). Since large variability between spontaneous beating frequencies has been reported (He et al., 2003; Ma et al., 2011), we created an additional rapid beating phenotype by scaling I_f_, I_NaCa_, and the SERCA-pump current (I_up_) by 2.5 (resulting AP biomarkers are shown in Table S6).

At 3D, cell injection was spatially modelled as in our previous study (Riebel et al., 2024). Briefly, six injection locations were chosen within the infarct and maintained for the small, medium, and large scar. The probability for an adult ventricular cell to be replaced by an hPSC-CM decreased with increasing distance from the injection location, assuming low migration of hPSC-CMs as observed experimentally in chronic scars (Poch et al., 2022). The hPSC-CM distribution transmurally and in the IZ versus BZ was calibrated to experimental measurements (Poch et al., 2022) – particularly, hPSC-CMs were more likely to settle in the subendocardium and in the IZ rather than the BZ. No hPSC-CMs were placed in the right ventricle or the RZ. For each scar, the spatial hPSC-CM distribution was maintained across simulations including from D0 to D14. Conductivities were calibrated to achieve a 10 cm/s conduction velocity in hPSC-CMs at D0 and maintained for all other hPSC-CM phenotypes (see Table S3). This resulted in an increased CV of 14.5 cm/s at D14, similar to experimental observations (Hansen et al., 2018).

### Identifying and optimising anti-arrhythmic ionic targets

To identify suitable anti-arrhythmic targets, a sensitivity analysis of the spontaneously beating (i.e., unpaced) Paci2020 model was performed as previously described (Riebel et al., 2021). Briefly, each ion channel’s conductance was scaled by +/-25% and the relative sensitivities on key AP biomarkers including upstroke velocity, peak membrane potential, APD_90_, maximum diastolic potential, and spontaneous beating frequency were computed as in Romero et al. (2009). The resulting relative sensitivities varied between −1 (strong negative correlation) and +1 (strong positive correlation).

Once targets to suppress automaticity were identified in the sensitivity analysis, we aimed to determine optimal combinations of ionic targets that also reduce re-entry susceptibility while preserving CaT_Amp_. For this, a population of 1,458 D0 and D14 hPSC-CM models was created by blocking I_f_ as well as upregulating G_K1_ and p_NaK_ in steps of 0.25 between 1 and 3 and G_Kr_ in steps of 0.5 between 1 and 5. Spontaneously beating models were discarded and in the remaining population, an optimisation index incorporating both safety (measured by APD_90_) and efficacy (measured by CaT_amp_) was calculated as:

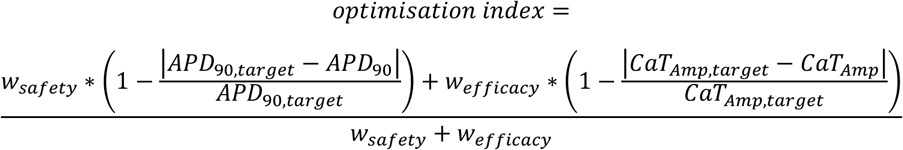

Here, the weights w_safety_ and w_efficacy_ were set to 1.5 and 1, respectively, to prioritise safety. Target APD_90_ can be used to create different hPSC-CM phenotypes; here we aimed towards adult values to reduce repolarisation gradients. Target CaT_Amp_ was always set to the maximum CaT_Amp_ achieved across the population (excluding any spontaneously beating models).

Modifying ionic current expression in hPSC-CMs has been achieved through gene editing techniques before injection (Marchiano et al., 2023). Hence, we combined optimisation indices at D0 and D14 for each ionic scaling combination, using also the maximum optimisation indices index_D0,max_ and index_D14,max_:

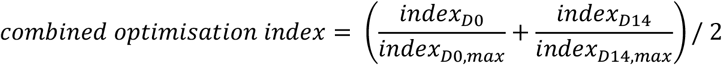

### Stimulation protocols to simulate sinus rhythm and quantify arrhythmic risk

The adult ventricular cardiomyocyte ToR-ORd model (Tomek et al., 2019), with RZ, BZ, or IZ remodelling applied (Zhou et al., 2024), and the Purkinje cell Trovato2020 model (Trovato et al., 2020) were paced at 1 Hz until steady state for 500 and 1,000 s, respectively, and then introduced into the biventricular model. As full coupling and therefore direct pacing of the hPSC-CMs in the ventricles could not be assumed, the hPSC-CM Paci2020 model (Paci et al., 2020), including D0, D14, rapid, and optimised phenotypes, were simulated under spontaneous (i.e., unpaced) conditions for 1,000 s before being introduced into the ventricles. The biventricular model was then paced for three 1 Hz beats applied at the top 25 elements of the bundle of His.

To assess automaticity-induced arrhythmias, biventricular models with either D0, rapid D0, D14, or rapid D14 hPSC-CMs were left without pacing for two seconds. If spontaneous beats occurred, ten 1 Hz beats were simulated to assess whether automaticity-induced arrhythmias would emerge under sinus pacing. To quantify re-entry burden, an S1-S2 protocol was employed: after the initial three 1 Hz beats (i.e., S1 stimuli), an S2 stimulus was applied in 15 different BZ locations between 350 to 570 ms after the S1. The different S2 locations were selected manually around the core of the small, medium, and large scars. Re-entry risk was measured as the VW (i.e., the range of S1-S2 intervals for which arrhythmias could be induced); considering S1-S2 intervals were increased in 20 ms increments, VWs were a multiple of 20.

This set up resulted in 1,458 single cell simulations to determine optimal ionic target combinations (9 scalings for GK1 * 9 scalings for pNaK * 9 scalings for GKr * 2 hPSC-CM phenotypes, i.e., D0 and D14) and 1,806 biventricular simulations (12 S1-S2 intervals * 15 S2 locations * (3 scars before cell injection + 3 scars at D0 + large scar at D14 + large scar with 3 (2 at D0, 1 at D14) optimised hPSC-CM models) + 6 simulations of spontaneous activity).

### Verification, validation, and uncertainty quantification

To establish trust in our modelling and simulation framework to mechanistically investigate cell therapy in chronic MI, we considered verification, validation, and uncertainty quantification principles (Musuamba et al., 2021; Viceconti, Pappalardo, et al., 2021). Verification of the monodomain solver was provided through benchmark tests (Berg et al., 2025; Sachetto Oliveira et al., 2018) and comparison to cellular model solvers (Riebel et al., 2024).

Validation of the cellular models was demonstrated through incorporation of biophysiological knowledge and experimental evidence of human cardiac cells in the original publications (Paci et al., 2020; Tomek et al., 2019; Trovato et al., 2020). Furthermore, predictive accuracy of these models has been established through *in silico* drug trials on contractility as well as arrhythmogenicity (Paci et al., 2021; Passini et al., 2021; Trovato et al., 2022, Trovato et al., 2025). Electrophysiological and structural remodelling post-MI was informed through experimental and clinical evidence as outlined above. Additionally, trust in the biventricular post-MI model was provided through comparison of simulated ECGs with clinical data (Riebel et al., 2024; Zhou et al., 2024).

In addition to previous sensitivity and convergence analyses (Riebel et al., 2024), we carried out further uncertainty quantification into the effects of steady state pacing duration and ionic concentrations on hPSC-CM APs at D0 and D14, as shown in Figures S8 and S9. These analyses showed small changes (less than 10%) in APD_90_.

## Data and Code Availability

Data and code will be made available upon publication.

## Acknowledgements

This project was funded by: a BBSRC PhD iCASE (BB/V509395/1) and Russell Studentship Agreement with AstraZeneca (R67719/CN001) for L.L.R. awarded to B.R., an Oxford-Bristol Myers Squibb Fellowship to X.Z. (R39207/CN063), a Wellcome Trust Senior Research Fellowship in Basic Biomedical Sciences to B.R. (214290/Z/18/Z), the Oxford BHF Centre of Research Excellence (RE/24/130024), and the EPSRC project CompBioMed X (EP/X019446/1).

The Paci2020 model was provided by Dr. Michelangelo Paci and the Computational Biophysics and Imaging Group (CBIG) at Tampere University Foundation.

An award for computer time was provided by the U.S. Department of Energy’s (DOE) Innovative and Novel Computational Impact on Theory and Experiment (INCITE) Program. This research used supporting resources at the Argonne Leadership Computing Facilities at Argonne National Laboratory, which is supported by the Office of Science of the U.S. DOE under Contract No. DE-AC02-06CH11357.

We would further like to thank Dr Max Cumberland for feedback and discussions.

Parts of this study were included in L.L. Riebel’s DPhil thesis at the University of Oxford, UK, titled Human-Based Multiscale Modelling and Simulation to Investigate Arrhythmias and Treatments in Ischemic Heart Disease (2024).

For the purpose of open access, the author has applied a Creative Commons Attribution (CC BY) public copyright licence to any Author Accepted Manuscript version arising from this submission.

## Author Contributions

L.L.R.: Conceptualisation, Methodology, Investigation, Writing – Original Draft, Z.J.W.: Conceptualisation, Methodology, Writing – Review & Editing, X.Z.: Methodology, Writing – Review & Editing, L.B.: Software, Writing – Review & Editing, C.T. & B.R.: Conceptualisation, Writing – Review & Editing, Supervision.

