## Supplemental Material for "*In Silico* Optimisation of Regenerative Cell Therapy in the Infarcted Human Ventricles to Mitigate Arrhythmic Burden"

#### S1. Supplemental Video Detail

Supplementary Video S1 shows automaticity-induced arrhythmias in the small (left) and large scar (right) at day 14 (D14) after virtual delivery of rapid stem cell-derived cardiomyocytes (hPSC-CMs, see Table S5 and Table S6 below). Note, that automaticity reaches a steady state at the end of the 10 second simulation in the small scar with a frequency of approximately 140 beats per minute, while in the large scar, automaticity causes re-entry and polymorphic tachycardia.

Supplementary Video S2 shows the only sustained re-entry at day 0 (D0) after cell delivery in the small scar. This was caused by stimulating 490 ms after the last sinus beat at S2 location 2 (basal septum). Note, that the re-entry is sustained through two Purkinje-myocyte junctions on the anterior wall.

Supplementary Video S3 depicts a re-entry before (left) and at D0 after (right) cell delivery in the large chronic scar, caused by stimulating 430 ms after the last sinus beat from S2 location 11 (apical). Note, that the re-entry in the control case lasts only for a single cycle, while the one at D0 after cell delivery is sustained.

Supplementary Video S4 compares a re-entry before (left) and after optimisation (right) of the cells at D14 in the large scar. Re-entry was induced by stimulating 410 ms after the last sinus beat at S2 location 2 (anteroseptal towards the base). Note, that the re-entry after delivery of the non-optimised cells is sustained, while after optimisation, it lasts only for a single cycle.

### S2. Supplemental Results

#### S2.1. Identifying anti-arrhythmic targets for hPSC-CMs

As shown in Figure S1 below, sensitivity analyses showed the D0 model's spontaneous beating rate most sensitive to pNaK (maximum sodium potassium pump current). Scaling pNaK by 0.7 and 0.85 was sufficient to abolish spontaneous activity in the D0 and D14 phenotype, respectively.

**Sensitivity analysis of unpaced hPSC-CMs**

| Parameter | MDP | $dV/dt_{max}$ | $V_{peak}$ | $APD_{90}$ | Frequency |
|---|---|---|---|---|---|
| $V_{max,up}$ | | 0.67 | | | 0.51 |
| $G_{CaL}$ | | 0.84 | 0.89 | 1.00 | |
| $k_{NaCa}$ | | | 0.26 | | 0.37 |
| $G_{NaF}$ | | 0.99 | | | |
| $G_{NaL}$ | | | | | |
| $P_{NaK}$ | | -1.00 | 1.00 | 0.25 | -1.00 |
| $G_{to}$ | | | | | |
| $G_{Kr}$ | | | -0.29 | -0.90 | |
| $G_{Ks}$ | | | | | |
| $G_{K1}$ | 1.00 | 0.40 | -0.41 | -0.74 | |
| $G_f$ | | -0.46 | | -0.29 | 0.35 |

Relative sensitivity < ±0.25 | Strong negative relation | Strong positive relation

*Figure S1. Sensitivity analysis of the spontaneous D0 (i.e., baseline Paci2020 (Paci et al., 2020)) hPSC-CM model to changes in ionic current conductances. Shown are relative sensitivities (1: strong positive relation, -1: strong negative relation), calculated as in Romero et al. (2009). Conductances were scaled by +/- 25% and hPSC-CMs left to beat spontaneously for 1,000 s.*

Figure S2, Figure S3, and Figure S4 below show ionic current and concentration traces underlying hPSC-CMs at D0 and D14 with and without scaling $p_{NaK}$, $G_{K1}$, and $G_f$.

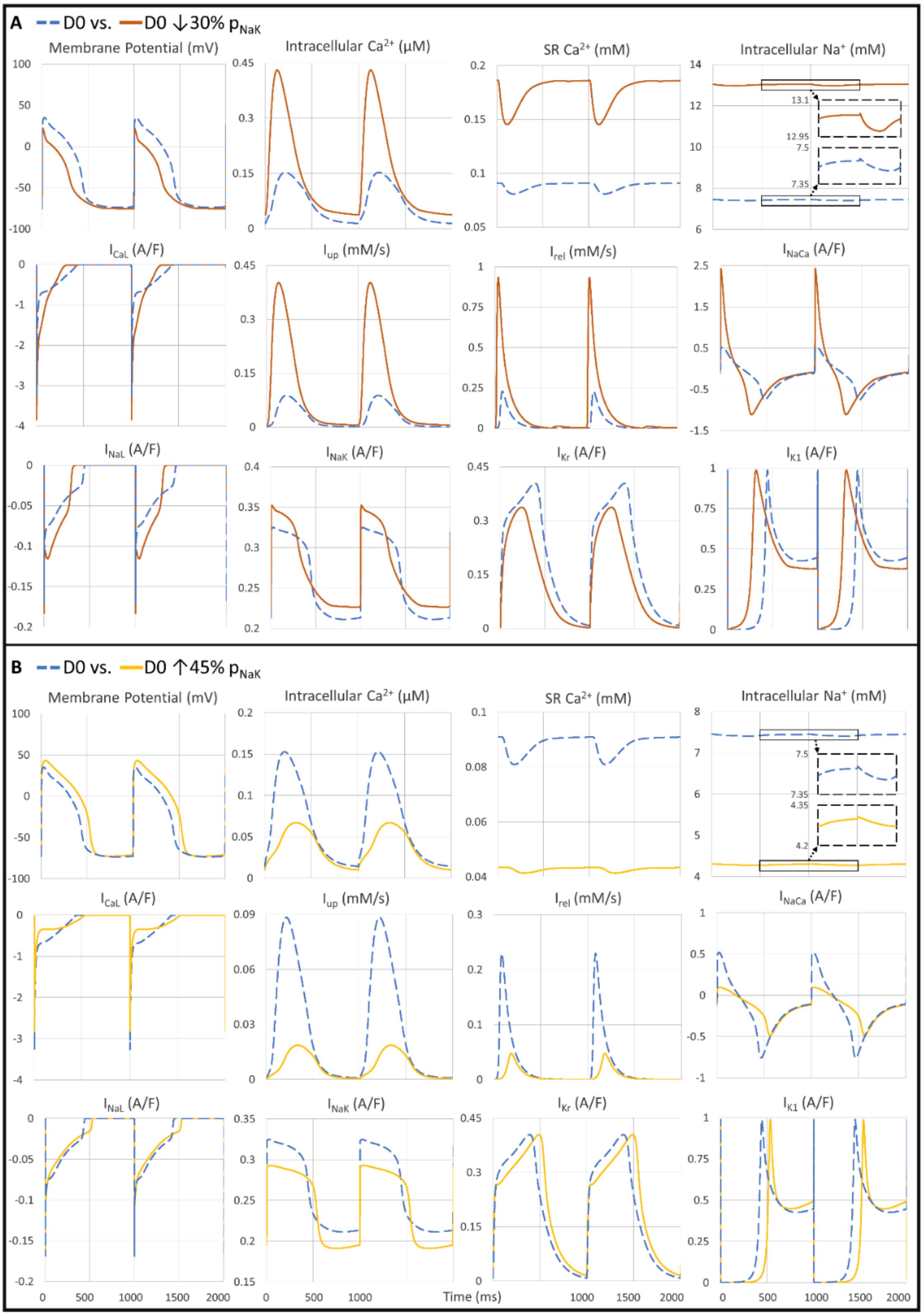

*Figure S2. Simulated action potential and ionic current traces of D0 hPSC-CMs in single cell after 1,000 paced 1 Hz beats. A) With and without downregulation of pNaK (maximum $Na^+$-$K^+$ pump current ($I_{NaK}$)). B) With and without upregulation of pNaK. Further shown are intracellular and SR $Ca^{2+}$ concentrations, intracellular $Na^+$ concentrations, $I_{CaL}$: L-type $Ca^{2+}$ current, $I_{up}$: SERCA pump current, $I_{rel}$: SR $Ca^{2+}$ release current, $I_{NaCa}$: $Na^+$-$Ca^{2+}$ exchanger current, $I_{NaL}$: late $Na^+$ current, $I_{Kr}$: rapid delayed outward $K^+$ current, and $I_{K1}$: inward rectifier $K^+$ current.*

Note, in the phenotype with reduced $p_{NaK}$ at D14 (Figure S3 A), the model had to be paced for 2000 rather than 1000 beats to reach steady state.

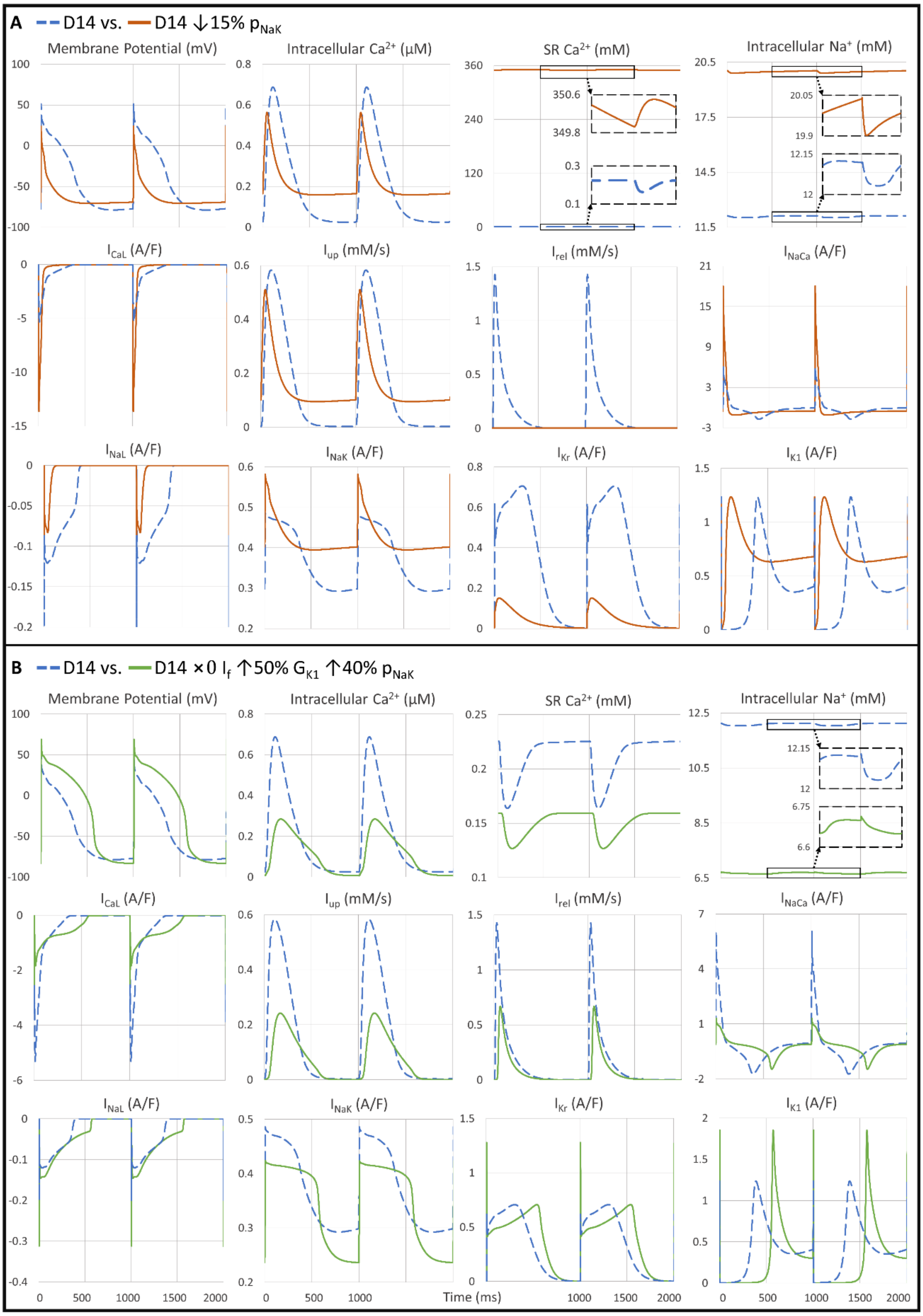

*Figure S3. Simulated action potential and ionic current traces of D14 hPSC-CMs in single cell after 1,000 paced 1 Hz beats. A) With and without downregulation of pNaK (maximum $Na^+$-$K^+$ pump current ($I_{NaK}$)). B) With and without upregulation of pNaK plus full block of the funny current ($I_f$) and upregulation of the inward rectifier $K^+$ current's ($I_{K1}$) conductance. Further shown are intracellular and SR $Ca^{2+}$ concentrations, intracellular $Na^+$ concentrations, $I_{CaL}$: L-type $Ca^{2+}$ current, $I_{up}$: SERCA pump current, $I_{rel}$: SR $Ca^{2+}$ release current (effectively 0 when reducing pNaK), $I_{NaCa}$: $Na^+$-$Ca^{2+}$ exchanger current, $I_{NaL}$: late $Na^+$ current, and $I_{Kr}$: rapid delayed outward $K^+$ current.*

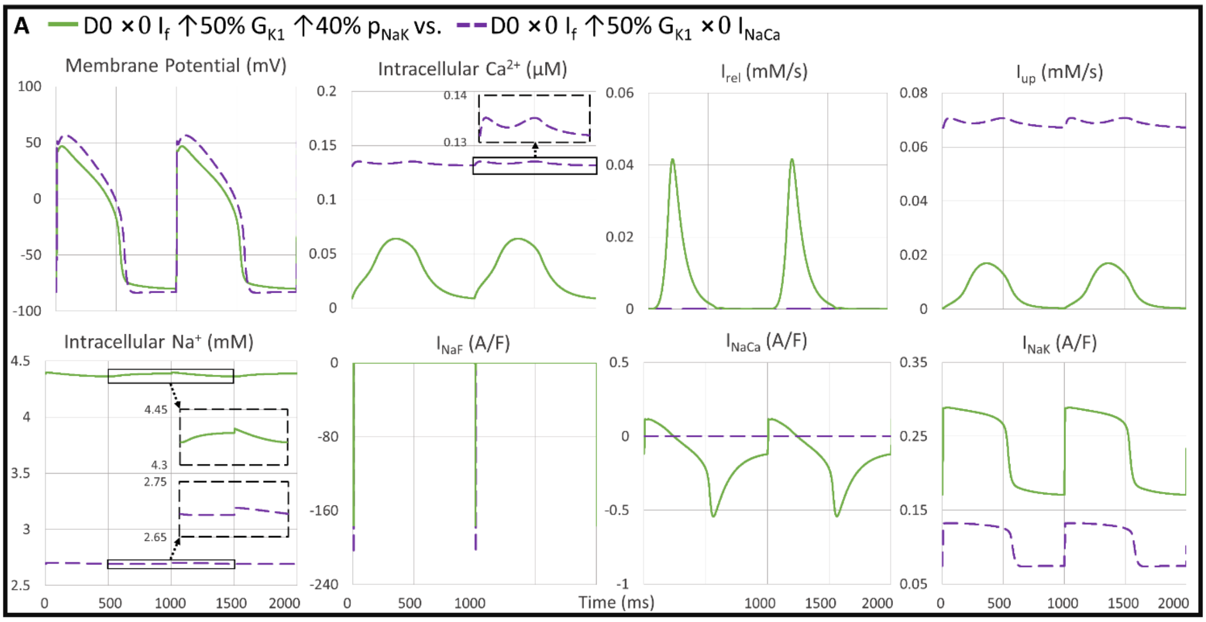

*Figure S4. Simulated action potential and ionic current traces after 1,000 paced 1 Hz beats of D0 hPSC-CMs with full block of the funny current ($I_f$) and upregulation of the inward rectifier $K^+$ current's ($I_{K1}$) conductance plus either A) upregulation of pNaK (maximum $Na^+$-$K^+$ pump current ($I_{NaK}$)) or B) full block of the $Na^+$-$Ca^{2+}$ exchanger current ($I_{NaCa}$, as suggested by Marchiano et al. (2023)). Further shown are intracellular $Ca^{2+}$ and $Na^+$ concentrations, $I_{rel}$: SR $Ca^{2+}$ release current (effectively 0 for $I_{NaCa}$ block), $I_{up}$: SERCA pump current, and $I_{NaF}$: fast $Na^+$ current.*

From the results presented in Figure S5, we chose the best performing ionic target combination. For Figure S6, we chose the ionic scaling combination with the third best result: fully blocking $I_f$ together with upregulating $G_{K1}$ by 1.25, $p_{NaK}$ by 1.75, and $G_{Kr}$ by 5, referred to in the text as "optimised" hPSC-CMs. We eliminated the top performing combination because it showed small beat-to-beat variations. We further discarded the second-best combination as it suggested that no $G_{K1}$ scaling was necessary, which may be specific to our virtual cell types. As the differences in $APD_{90}$ and $CaT_{Amp}$ between the second and third-best options were only 2 ms and 0.001 µM, respectively, we finally chose to proceed with the third option that maintained $G_{K1}$ scaling at 1.25. The optimised cells at D0 and D14 had an $APD_{90}$ of 169 and 230 ms, respectively.

| GK1 | GKr \ PNaK | 1.5 | 1.75 | 2 | 2.25 | 2.5 | 2.75 | 3 |
|---|---|---|---|---|---|---|---|---|
| 1 | 1 | | | | | | | |
| | 1.5 | | | | | | | |
| | 2 | | | | | | | |
| | 2.5 | | 0.86 | 0.84 | 0.83 | 0.81 | 0.81 | 0.80 |
| | 3 | | 0.872 | 0.86 | 0.85 | 0.84 | 0.84 | 0.83 |
| | 3.5 | | 0.85 | 0.84 | 0.84 | 0.83 | 0.82 | 0.82 |
| | 4 | | 0.80 | 0.79 | 0.78 | 0.78 | 0.77 | 0.77 |
| | 4.5 | 0.77 | 0.75 | | | | | |
| | 5 | | | | | | | |
| 1.25 | 1 | | | | | | | |
| | 1.5 | | | | | | | |
| | 2 | | | | | | | |
| | 2.5 | | 0.86 | 0.83 | 0.82 | 0.81 | 0.80 | 0.80 |
| | 3 | | 0.872 | 0.86 | 0.85 | 0.84 | 0.84 | 0.83 |
| | 3.5 | | 0.86 | 0.84 | 0.83 | 0.83 | 0.83 | 0.82 |
| | 4 | | 0.80 | 0.79 | 0.78 | 0.78 | 0.77 | 0.77 |
| | 4.5 | | 0.75 | | | | | |
| | 5 | | | | | | | |
| 1.5 | 1 | | | | | | | |
| | 1.5 | | | | | | | |
| | 2 | 0.79 | | | | | | |
| | 2.5 | | 0.86 | 0.84 | 0.82 | 0.81 | 0.80 | 0.79 |
| | 3 | | **0.874** | 0.86 | 0.85 | 0.84 | 0.84 | 0.83 |
| | 3.5 | | 0.86 | 0.84 | 0.84 | 0.83 | 0.83 | 0.82 |
| | 4 | | | 0.79 | 0.78 | 0.78 | 0.78 | 0.77 |
| | 4.5 | | | 0.75 | | | | |
| | 5 | | | | | | | |
| 1.75 | 1 | | | | | | | |
| | 1.5 | | | | | | | |
| | 2 | 0.80 | 0.75 | | | | | |
| | 2.5 | | 0.86 | 0.84 | 0.82 | 0.81 | 0.80 | 0.79 |
| | 3 | | | 0.86 | 0.85 | 0.84 | 0.84 | 0.84 |
| | 3.5 | | | 0.84 | 0.84 | 0.83 | 0.83 | 0.83 |
| | 4 | | | | 0.79 | 0.78 | 0.78 | 0.77 |
| | 4.5 | | | | | | | |
| | 5 | | | | | | | |
| 2 | 1 | | | | | | | |
| | 1.5 | | | | | | | |
| | 2 | | 0.75 | | | | | |
| | 2.5 | | | 0.84 | 0.82 | 0.81 | 0.80 | 0.79 |
| | 3 | | | | 0.85 | 0.84 | 0.84 | 0.84 |
| | 3.5 | | | | | 0.83 | 0.83 | 0.83 |
| | 4 | | | | | 0.78 | 0.78 | 0.77 |
| | 4.5 | | | | | | | |
| | 5 | | | | | | | |
| 2.25 | 1 | | | | | | | |
| | 1.5 | | | | | | | |
| | 2 | | | | | | | |
| | 2.5 | | | | | | | |
| | 3 | | | | | | | |
| | 3.5 | | | | | | | |
| | 4 | | | | | | | |
| | 4.5 | | | | | | | |
| | 5 | | | | | | | |
| 2.5 | 1 | | | | | | | |
| | 1.5 | | | | | | | |
| | 2 | | | | | | | |
| | 2.5 | | | | | | | |
| | 3 | | | | | | | |
| | 3.5 | | | | | | | |
| | 4 | | | | | | | |
| | 4.5 | | | | | | | |
| | 5 | | | | | | | |

*Figure S5. Optimisation index with a target $APD_{90}$ of 330 ms, computed for full block of the funny current ($I_f$) and different combinations of upscaling the inward rectifier $K^+$ current conductance ($G_{K1}$), the maximum $Na^+$-$K^+$ pump current ($P_{NaK}$), and the rapid delayed outward rectifier $K^+$ current conductance ($G_{Kr}$). Dark green indicates a high scoring. Scores below 0.75 are not numbered. Note, no score was computed if the cells were still beating spontaneously or could not be paced. Highlighted is the best performing scaling combination.*

| | PNaK | 1.5 | 1.75 | 2 | 2.25 | 2.5 | 2.75 | 3 |
|---|---|---|---|---|---|---|---|---|
| GK1 | GKr | | | | | | | |
| | 1 | | | | | | | |
| | 1.5 | | | | | | | |
| | 2 | | | | | | | |
| | 2.5 | | | | | | | |
| 1 | 3 | | | | | | | |
| | 3.5 | | | | | | | |
| | 4 | | | | | | | |
| | 4.5 | 0.87 | 0.84 | 0.82 | 0.81 | 0.80 | 0.79 | 0.79 |
| | 5 | 0.95 | 0.9270 | 0.92 | 0.91 | 0.90 | 0.89 | 0.89 |
| | 1 | | | | | | | |
| | 1.5 | | | | | | | |
| | 2 | | | | | | | |
| | 2.5 | | | | | | | |
| 1.25 | 3 | | | | | | | |
| | 3.5 | | | | | | | |
| | 4 | | | | | | | |
| | 4.5 | | 0.83 | 0.81 | 0.80 | 0.79 | 0.78 | 0.77 |
| | 5 | | 0.9266 | 0.91 | 0.90 | 0.89 | 0.88 | 0.87 |
| | 1 | | | | | | | |
| | 1.5 | | | | | | | |
| | 2 | | | | | | | |
| | 2.5 | | | | | | | |
| 1.5 | 3 | | | | | | | |
| | 3.5 | | | | | | | |
| | 4 | | | | | | | |
| | 4.5 | | | 0.81 | 0.79 | 0.78 | 0.77 | 0.76 |
| | 5 | | | 0.90 | 0.89 | 0.88 | 0.87 | 0.87 |
| | 1 | | | | | | | |
| | 1.5 | | | | | | | |
| | 2 | | | | | | | |
| | 2.5 | | | | | | | |
| 1.75 | 3 | | | | | | | |
| | 3.5 | | | | | | | |
| | 4 | | | | | | | |
| | 4.5 | | | | 0.79 | 0.78 | 0.77 | 0.76 |
| | 5 | | | | 0.89 | 0.88 | 0.87 | 0.86 |
| | 1 | | | | | | | |
| | 1.5 | | | | | | | |
| | 2 | | | | | | | |
| | 2.5 | | | | | | | |
| 2 | 3 | | | | | | | |
| | 3.5 | | | | | | | |
| | 4 | | | | | | | |
| | 4.5 | | | | | | 0.77 | 0.76 |
| | 5 | | | | | | | 0.86 |
| | 1 | | | | | | | |
| | 1.5 | | | | | | | |
| | 2 | | | | | | | |
| | 2.5 | | | | | | | |
| 2.25 | 3 | | | | | | | |
| | 3.5 | | | | | | | |
| | 4 | | | | | | | |
| | 4.5 | | | | | | | |
| | 5 | | | | | | | |
| | 1 | | | | | | | |
| | 1.5 | | | | | | | |
| | 2 | | | | | | | |
| | 2.5 | | | | | | | |
| 2.5 | 3 | | | | | | | |
| | 3.5 | | | | | | | |
| | 4 | | | | | | | |
| | 4.5 | | | | | | | |
| | 5 | | | | | | | |

*Figure S6. Optimisation index with a target $APD_{90}$ of 170 ms, computed for full block of the funny current ($I_f$) and different combinations of upscaling the inward rectifier $K^+$ current conductance ($G_{K1}$), the maximum $Na^+$-$K^+$ pump current ($p_{NaK}$), and the rapid delayed outward rectifier $K^+$ current conductance ($G_{Kr}$). Dark green indicates a high scoring. Scores below 0.75 are not numbered. Note, no score was computed if the cells were still beating spontaneously or could not be paced. Highlighted are the top three performing scaling combinations with the selected one in yellow.*

### S3. Supplemental Methods

#### S3.1. Human electrophysiological modelling and simulation of chronic MI

Table S1 below summarises ionic scaling factors that were applied to model epicardial (based on the baseline endocardial) human ventricular cardiomyocytes and an apicobasal gradient.

*Table S1. Ionic current scaling factors to produce I) epicardial human ventricular cardiomyocytes from the baseline endocardial phenotype as in [1]Tomek et al. (2019, 2020) resulting in a transmural action potential duration gradient of 35 ms, and II) a ventricular apex to base gradient as established by [2]Okada et al. (2011) and Mincholé et al. (2019) resulting in an apicobasal repolarisation gradient of 10 ms. $V_m$ denotes the membrane voltage.*

| Model parameter | Epicardial cell type[1] | Apex to base gradient[2] |
|---|---|---|
| $G_{NaL}$ | *0.6 | |
| $P_{NaK}$ | *0.9 | |
| $G_{to}$ | *2.0 | |
| $delta_{epi}$ | $= 1 - \frac{0.95}{1 + e^{\frac{V_m+70}{5}}}$ | |
| $G_{Kr}$ | *1.3 | |
| $G_{Ks}$ | *1.4 | *gradient between 0.2 at the base and 5.0 at the apex |
| $G_{K1}$ | *1.2 | |
| $G_{Kb}$ | *0.6 | |
| $G_{NaCa}$ | *1.1 | |
| $P_{Ca}$ | *1.2 | |
| $J_{upnp}$ | *1.3 | |
| $J_{upp}$ | *1.3 | |
| $cmdn_{max}$ | *1.3 | |

Table S2 below outlines remodelling of cellular ion channel conductances and time constants that was applied to simulate chronically infarcted human cardiomyocytes in the infarct, border, and remote zone.

*Table S2. Border and remote zone remodelling as established by Zhou et al. (2024). Infarct zone modelling as suggested by [7]Ringenberg et al. (2014). Supporting experimental data from [1]Valdivia et al. (2005), [2]Hegyi et al. (2018), [3]Jiang et al. (2002), [4]Høydal et al. (2018), [5]Maier et al. (2003), and [6]Hoch et al. (1999).*

| Model Parameter | Remote Zone | Border Zone | Infarct Zone |
|---|---|---|---|
| $V_m$ (set not scaled) | | | -60 mV[7] |
| $G_{Na}$ | 0.43[1] | 0.43[1] | |
| $G_{NaL}$ | 1.413[2] | 1.275[2] | |
| $G_{Kr}$ | 0.87[2] | 0.89[2] | |
| $G_{K1}$ | | 0.76[2] | |
| $G_{KCa}$ | 2[2] | 2[2] | |
| $P_{Ca}$ | | 0.7[2] | |
| $J_{up}$ | 0.4[3,4] | 0.4[3,4] | |
| $tau_{relp}$ | 6[5] | 6[5] | |
| aCaMK | 1.5[6] | 1.5[6] | |
| $G_{ClCa}$ | 1.25[2] | 1.25[2] | |

#### S3.2. Modelling and simulation of cell therapy

Table S3 below shows conduction velocities that were achieved in the adult myocardium as well as the virtually injected hPSC-CMs.

*Table S3. Conduction velocities of adult ventricular tissue and hPSC-CMs. Using conductivity tensors of (2.60, 1.05, 1.60) mS/cm in the remote zone, (0.60, 0.35, 0.45) mS/cm in the border and infarct zone, and (0.2, 0.2, 0.2) mS/cm in hPSC-CMs. Measured in a 20x7x3 cm tissue slab stimulated in the bottom corner, as suggested in benchmark tests by Niederer et al. (2011). Note, hPSC-CM connectivity was modelled as isotropic.*

| **Cell type** | | **Conduction Velocity (cm/s)** | | |
|---|---|---|---|---|
| | | Fibre | Sheet | Normal |
| Adult ventricular myocardium | Remote Zone | 49 | 27 | 34 |
| | Border Zone | 19 | 12 | 15 |
| | Infarct Zone | 9 | 7 | 8 |
| Stem cell-derived cardiomyocytes | D0 (baseline Paci2020 model) | 10 | | |
| | Rapid D0 | 9 | | |
| | D0 with ×0 $I_f$ ↑50% $G_{K1}$ ↑75% $p_{NaK}$ ↑200% $G_{Kr}$ | 12.5 | | |
| | D0 with ×0 $I_f$ ↑25% $G_{K1}$ ↑75% $p_{NaK}$ ↑400% $G_{Kr}$ | 12.5 | | |
| | D14 | 14.5 | | |
| | Rapid D14 | 12.5 | | |
| | D14 with ×0 $I_f$ ↑25% $G_{K1}$ ↑75% $p_{NaK}$ ↑400% $G_{Kr}$ | 18 | | |

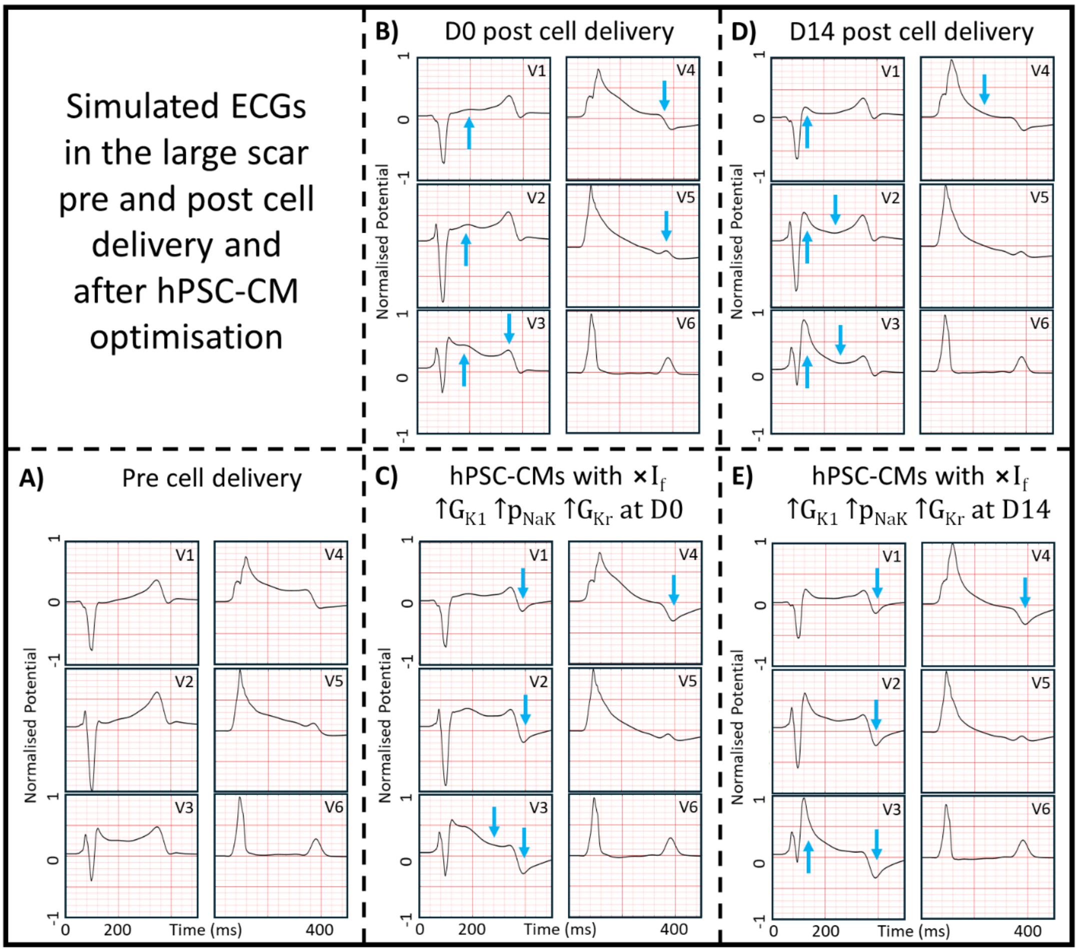

*Figure S7. Simulated ECG before and at D0 and D14 after virtual delivery of stem cell-derived cardiomyocytes (hPSC-CMs) before and after optimisation (i.e., $I_f$ block and $G_{K1}$, $p_{NaK}$, and $G_{Kr}$ upregulation). Blue arrows indicate prominent changes from panels A) to B), B) to C), B) to D), and D) to E).*

Next, simulated ECGs from before and after virtual cell injection are shown in Figure S7 above. Delivery of hPSC-CMs at D0 caused ST-elevation in leads V1 to V3 as well as T-wave depression in leads V4 and V5. Virtual injection of hPSC-CMs at D14 increased QRS fragmentation and J-point elevation in leads V1 to V3 as well as depressed T-waves in leads V4 and V5. Furthermore, as conduction velocity increased in hPSC-CMs from D0 to D14 (see Table S3), activation of the scar improved, which is visible in the ECG as reduced QRS-prolongation. Delivery of optimised hPSC-CMs at D0 and D14 further depressed the T-wave, also in leads V2 and V3.

Table S4 below details input parameters for the Monodomain solver MonoAlg3D (Berg et al., 2025; Sachetto Oliveira et al., 2018).

*Table S4. Parameter files to configure MonoAlg3D (Berg et al., 2025; Sachetto Oliveira et al., 2018) for 3D simulations.*

<table>
<tr><th>MonoAlg3D Parameter</th><th>ToR-ORd</th><th>Paci2020</th><th>Trovato</th></tr>
<tr><td>Space adaptivity</td><td colspan="3">false</td></tr>
<tr><td>PDE timestep</td><td colspan="3">0.01</td></tr>
<tr><td>PDE solved in the…</td><td colspan="3">GPU</td></tr>
<tr><td>ODE time step</td><td colspan="3">0.01</td></tr>
<tr><td>ODE solver method</td><td colspan="3">Rush-Larsen + Euler</td></tr>
<tr><td>ODE adaptivity</td><td colspan="3">false</td></tr>
<tr><td>ODE solved in the…</td><td colspan="2">GPU</td><td>CPU</td></tr>
<tr><td>Linear system solver gradient function</td><td colspan="2">Biconjugate</td><td>Conjugate</td></tr>
<tr><td>Linear system solver tolerance</td><td colspan="2">1e-12</td><td>1e-16</td></tr>
</table>

Table S5 and Table S6 below introduce scaling factors for ionic current conductances to simulate baseline and rapid hPSC-CM phenotypes at D0 and D14 and further show their resulting action potential biomarkers.

*Table S5. Ion channel conductance scaling factors to produce D14 (based on experimental gene expression data from Marchiano et al. (2023, Figure 1F)) and rapid beating hPSC-CMs using Paci2020 (Paci et al., 2020) as the baseline D0 model.*

| Model Parameter | D14 | Rapid D0 | Rapid D14 |
|---|---|---|---|
| $G_f$ | 1.25 | 2.50 | 3.125 |
| $G_{K1}$ | 1.25 | | 1.250 |
| $G_{to}$ | 1.50 | | 1.500 |
| $G_{Ks}$ | 2.00 | | 2.000 |
| $G_{Kr}$ | 1.75 | | 1.750 |
| $G_{NaF}$ | 2.50 | | 2.500 |
| $k_{NaCa}$ | 2.00 | 2.50 | 5.000 |
| $G_{CaL}$ | 3.00 | | 3.000 |
| $V_{max,up}$ | | 2.50 | 2.500 |

*Table S6. AP biomarkers for D0, D14, and rapid hPSC-CM phenotypes after 1,000 seconds of spontaneous activity using Paci2020 (Paci et al., 2020). DDR: diastolic depolarisation rate, MDP: maximum diastolic potential, $dV/dt_{max}$: maximum upstroke velocity, $V_{m,peak}$: peak $V_m$, $APD_{90}$: APD at 90% repolarisation, F: spontaneous beating frequency..*

| hPSC-CM phenotype | DDR (V/s) | MDP (mV) | $dV/dt_{max}$ (V/s) | $V_{m,peak}$ (mV) | $APD_{90}$ (mV) | F (bpm) |
|---|---|---|---|---|---|---|

| | | | | | | |
|---|---|---|---|---|---|---|
| D0 | 0.01 | -74.87 | 20.54 | 27.14 | 402.42 | 35 |
| Rapid D0 | 0.02 | -74.90 | 26.49 | 18.76 | 424.27 | 52 |
| D14 | 0.02 | -78.79 | 172.26 | 36.53 | 446.31 | 47 |
| Rapid D14 | 0.03 | -78.57 | 168.01 | 29.94 | 414.57 | 66 |

#### S3.3. Verification, validation, and uncertainty quantification

To quantify uncertainty, we carried out further sensitivity analyses. First, as mentioned in the stimulation protocols in the main text, biventricular simulations used the unpaced cellular hPSC-CM models after 1,000 as a starting state. We chose unpaced rather than paced activity, as coupling between hPSC-CMs and surrounding tissue, particularly within the scar, may be incomplete impacting their pacing. However, in single cell, the optimised hPSC-CMs' spontaneous beating was suppressed and only a few APs occurred at the start of the unpaced 1,000 s. This leads to uncertainty regarding the cells' phenotype after possible pacing in the ventricles, especially as we changed $I_{NaK}$, which may substantially affect ionic concentrations. Hence, we compared the APs of optimised hPSC-CMs (full $I_f$ block and 25%, 75%, and 400% $I_{K1}$ $I_{NaK}$, and $I_{Kr}$ upregulation, respectively) after 1,000 s of unpaced activity followed by 3 paced beats (mimicking the biventricular simulation setting) versus 1,000 paced beats. As shown in Figure S8 A) below, this produced small changes in the D0 phenotype ($APD_{90}$ difference of approximately 10 ms). At D14 (panel B), the $CaT_{Amp}$ increased about 3-fold over time and the AP shape changed more substantially (reduced AP peak and steeper repolarisation). However, $APD_{90}$ changed only from 230 to 245 ms with increased pacing duration.

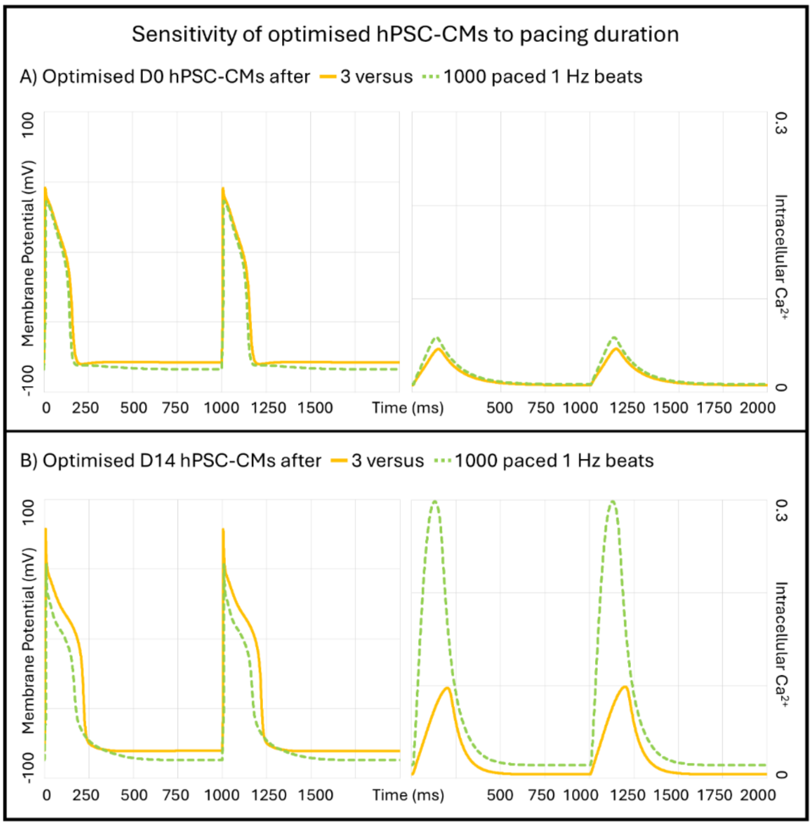

*Figure S8. Sensitivity of optimised stem cell-derived cardiomyocytes (hPSC-CMs) to pacing duration. hPSC-CMs at A) D0 and B) D14 with full block of the funny current, 25% upregulation of the inward rectifier $K^+$ current's conudctance, 75%*

*upregulation of maximum Na⁺-K⁺ pump current, and 400% upregulation of the rapid delayed outward rectifier K⁺ current' conductance. Simulated in single cell for 1,000 s of unpaced activity followed by 3 s (solid yellow traces) or 1,000 s (dashed green traces) of 1 Hz pacing.*

Next, we investigated the sensitivity to differences in ionic concentrations, as these differ slightly in the hPSC-CM Paci2020 model and the adult ventricular cardiomyocyte ToR-ORd model. Specifically, extracellular $K^+$ and $Na^+$ concentrations are 151 versus 140 and 5.4 versus 5.0 mM in the Paci2020 versus ToR-ORd model, respectively. Considering that ionic concentrations are variable spatially and over time, in our simulations, we did not change these, also allowing for comparability to previous studies. Comparison on the effect of changing the Paci2020 concentrations to the ToR-ORd values, shown in Figure S9, showed small changes in the spontaneous beating frequency (35 bpm before versus 36 bpm after changing concentrations) and the paced $APD_{90}$ (455 before versus 440 ms after changing concentrations).

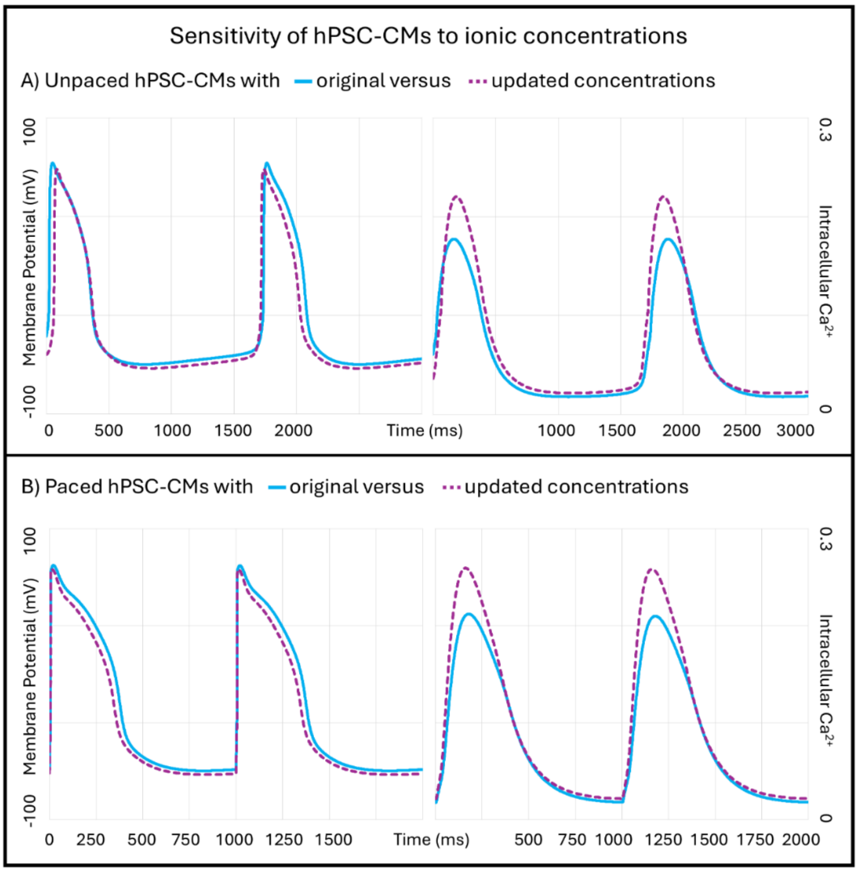

*Figure S9. Sensitivity of single cell hPSC-CMs at D0 to ionic concentrations. Original hPSC-CM Paci2020 model (Paci et al., 2020) with extracellular Na⁺ of 151 mM and extracellular K⁺ of 5.4 mM (solid blue traces) or using ionic concentrations from the adult ventricular ToR-ORd model (Tomek et al., 2019, 2020), i.e., extracellular Na⁺ of 140 mM and extracellular K⁺ of 5 mM (dashed purple traces). A) 1,000 s of spontaneous beating were simulated and B) followed by 3 s of 1 Hz pacing.*
